# One species, two genomes: A critical assessment of inter-isolate variation and identification of assembly incongruence in *Haemonchus contortus*

**DOI:** 10.1101/384008

**Authors:** Jeff A. Wintersinger, Grace M. Mariene, James D. Wasmuth

## Abstract

**Background:** Numerous quality issues may compromise genomic data’s representation of its underlying organism. In this study, we compared two genomes published by different research groups for the parasitic nematode *Haemonchus contortus*, corresponding to divergent isolates. We analyzed differences between the genomes, attempting to ascertain which were attributable to legitimate biological differences, and which to technical error in one or both genomes.

**Results:** We found discrepancies between the *H. contortus* genomes in both assembly and annotation. The genomes differed in representation of genes that are highly conserved across eukaryotes, with clear evidence of misassembly underlying conserved genes missing from one genome or the other. Only 45% of genes in one genome were orthologous to genes in the other genome, with one genome exhibiting almost as much orthology to *C. elegans* as its counterpart *H. contortus* strain. The two genomes differed substantially in probable causes underlying this unexpectedly low orthology. One genome included many more inparalogues than the other, and more frequently assembled inparalogues together on the same portions of contiguous sequence. It also exhibited cases of better-conserved gene position relative to *C. elegans*.

**Conclusion:** The discrepancies between the two genomes far exceeded those expected as a consequence of biological differences between the two *H. contortus* isolates. This implies substantial quality issues in one or both genomes, suggesting that researchers must exercise caution when using genomic data for newly sequenced species.

## Background

In the decade following publication of the human genome, the use of genomic sequence data has increased dramatically [1], with a continued exponential drop in sequencing costs pushing researchers to utilize sequencing data in sundry studies spanning biology and medicine.

Genomic data production, however, is complicated, suffering from numerous issues that compromise the resulting data’s reflection of its underlying organism. The precision of genomic data can easily be mistaken for accuracy—though every nucleotide, gene boundary, and splice site is denoted exactly, uncertainty in the processes producing them is poorly represented in the resulting genome [2]. Thus, researchers who draw on genomic data without understanding their potential pitfalls will place undue faith in the validity of results built atop genomic foundations.

Though sequencing costs have dropped precipitously, assembly, annotation, and analysis remain complex processes [3] subject to problems affecting the quality of the resulting genome [4]. The short-read sequencing technology prevalent today struggles to resolve repetitive regions [5], resulting in repeat sequences being collapsed, or physically distant regions being mistakenly adjoined because they flank highly similar sequences. With sequencing platforms being 85.0% to 99.9% accurate in each called base [4], distinguishing misread bases from legitimate variation stemming from rare alleles is difficult. This problem is compounded when dealing with samples drawn from pools of multiple individuals, as such samples may contain considerably more than the two alleles potentially present in a single diploid organism.

Genome annotation—i.e., the construction of gene models—is no less arduous. While computational methods of *ab initio* gene prediction achieve accuracy approaching 100% given sufficient training [6], this process requires numerous existing high-quality gene models specific to the studied organism, which are typically unavailable for newly sequenced species. Moreover, these methods accurately denote intron/exon boundaries within genes in only 60% to 70% of cases [6]. In addition, while transcriptomic data yields high-confidence coding sequences that can be used for annotation, it is hardly comprehensive—it will capture only transcribed genes, the set of which will change according to environmental conditions and the organism’s life-cycle stage.

Molecular biologists who use genome sequences suffer from a pervasive lack of awareness concerning assembly and annotation issues that compromise genomic data’s representation of its underlying organism, potentially leading to erroneous biological interpretations. Thus, it is critical to promote understanding of these problems, both amongst researchers drawing on genomic data, and amongst labs producing such data who are unseasoned in these tasks. In this study, we compared two recently published genomes for the Clade V parasitic nematode *Haemonchus contortus.* One genome corresponds to the laboratory isolate MHco3 derived from an African *H. contortus* sample (herein referred to as the MHco3 genome) [7], and the other to an Australian field isolate (herein referred to as the McMaster genome) [8]. By investigating discrepancies in the genomes’ assemblies and annotations, we aimed to infer which resulted from technical error, and which from legitimate biological variation. The methods we present here may now be applied to other species for which multiple genomes exist.

Past studies comparing genomic data sets have examined multiple annotations available for a single genome assembly [9–11]. Our work is novel in that it also accounts for assembly variation, introducing significant complexity because of assembly’s influence on annotation quality. Reannotating an improved *Bos taurus* assembly, for example, yielded substantial changes in 40% of genes, with 660 protein-coding genes lost in the improved version [12]. While other studies have compared multiple isolates of the same organism, they have been limited in scope. An examination of two isolates of the fungus *Aspergillus oryzae* deemed differences to be legitimate biological variations, without considering which may be attributable to technical error [13]. Conversely, a study of the SRS gene family across three strains of the parasitic protozoan *Toxoplasma* gondii proposed that large chromosomal rearrangements are likely indicative of assembly error rather than true biological variation [14]. Our *H. contortus* analysis expands these efforts, examining interisolate variation on a genome-wide scale while encompassing technical error as a potential source of interisolate discrepancies. This study also accounts for greater genomic size compared to past work. Relative to the 36 Mb *A. oryzae* [13] and 65 Mb *T. gondii* [14] genomes, the *H. contortus* assemblies are between 320-380 Mb, listing twice as many genes (Table 1).

**Table 1:**
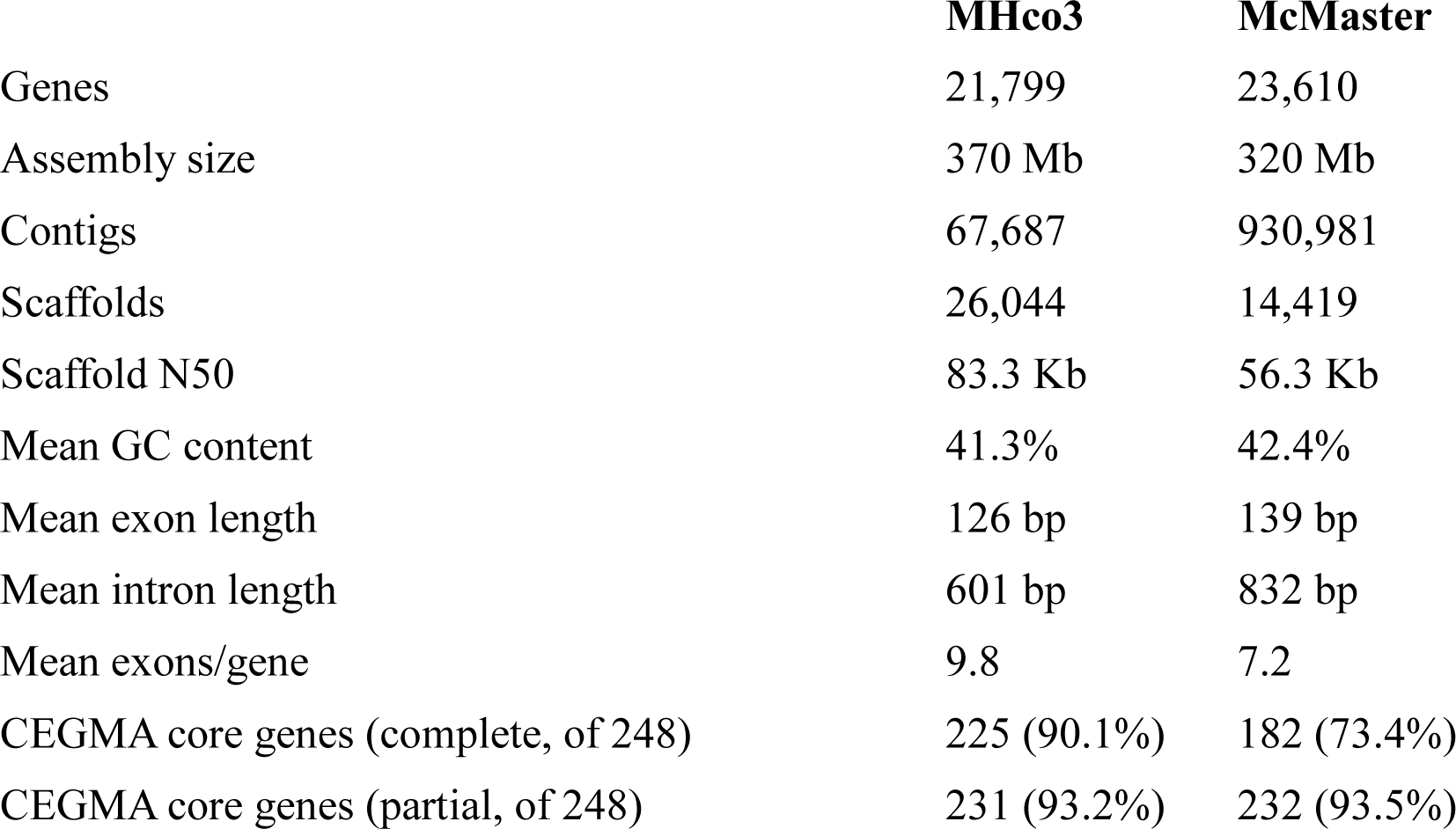
Comparison of metrics characterizing the MHco3 [7] and McMaster [8] genomes included in their respective publications.

|  | **MHco3** | **McMaster** |
| --- | --- | --- |
| Genes | 21,799 | 23,610 |
| Assembly size | 370 Mb | 320 Mb |
| Contigs | 67,687 | 930,981 |
| Scaffolds | 26,044 | 14,419 |
| Scaffold N50 | 83.3 Kb | 56.3 Kb |
| Mean GC content | 41.3% | 42.4% |
| Mean exon length | 126 bp | 139 bp |
| Mean intron length | 601 bp | 832 bp |
| Mean exons/gene | 9.8 | 7.2 |
| CEGMA core genes (complete, of 248) | 225 (90.1%) | 182 (73.4%) |
| CEGMA core genes (partial, of 248) | 231 (93.2%) | 232 (93.5%) |

*H. contortus* is an economically important parasite that has infected hundreds of millions of sheep and goats globally, producing losses of tens of billions of dollars per annum [15, 16]. Feeding on capillary blood in stomach mucosa [16], *H. contortus* causes anemia and hemorrhagic gastritis in infected animals. Infections on farms spread rapidly, with a single female parasite producing up to 10,000 eggs per day, and some hosts being infected with thousands of worms [17]. Though *H. contortus* infections are commonly treated via anthelmintic drugs, the worm has demonstrated the ability to rapidly develop resistance [18]. As *in vivo H. contortus* studies, including genetic crosses [19], can be undertaken in the parasite’s natural host, *H. contortus* serves as an ideal model for strongylid drug resistance and vaccine development [7].

*Haemonchus’* suitability as a study subject extends beyond its economic importance and use as a model of drug resistance [15, 16, 18]. *H. contortus* presents an ideal opportunity for genomic comparison, as the two concurrently published genomes considered here each utilized state-of-the-art sequencing, assembly, and annotation methods [7, 8]. Thus, the resulting data sets should represent their corresponding isolates well. Nevertheless, as both projects used different technologies and methods, discrepancies between the data sets may correspond to instances of one genome’s method faltering where the other succeeded. Additionally, with *H. contortus* exhibiting substantial genomic diversity between isolates due to its large effective population size [20, 21], we must also account for considerable biologically valid interisolate variation. The resulting methods can thus be applied to other eukaryotic genomes whose size and complexity are comparable to that of *H. contortus*.

The metrics published alongside each genome demonstrate significant discord between the two (Table 1). MHco3 is 370 Mb long relative to McMaster’s 320 Mb, while McMaster has 1811 more annotated genes than MHco3. Though lower or higher values for assembly length or gene count are not reflective of superior accuracy, the magnitude of intergenome differences for both values exceeds that which we would expect from legitimate biological variation, suggesting that some of this variance is attributable to technical error in one or both genomes. Our study capitalizes on these discrepancies, exploring both their extent and implications.

Entering this study, our hypothesis was that minimal legitimate variation existed between the isolates, and thus that most intergenome disparities would stem from technical error. Legitimate variations would likely be limited to duplications and deletions of non-critical genes, as well as small-scale rearrangements. The only existing metric permitting quality comparisons was scaffold N50, with the N50 in MHco3 being 83.3 Kb relative to 56.3 Kb in McMaster, suggesting MHco3’s assembly is more contiguous and thus more accurate than McMaster’s. This observation comes with the caveat, however, that N50 does not necessarily reflect quality. A higher N50 may instead indicate a less stringent assembly process that erroneously adjoined non-contiguous sequences. While our results indicate that MHco3 likely better represents *H. contortus* than McMaster, they also show pervasive quality issues in each genome, and much greater intergenome variation than expected. These issues make difficult the task of identifying legitimate biological variations between the two strains.

## Results

### Numerous CEGs are missing from each assembly

Of the 248 most highly conserved CEGs, 166 (67%) could be recovered as complete copies in both genomes (Figure 1A), while 209 (84%) could be found in complete or partial forms (Figure 1B). If we consider each genome individually, MHco3 fared better than McMaster—50 CEGs present completely in MHco3 were found only as partial copies or were absent altogether in McMaster (Figure 1A), while only seven CEGs were complete in McMaster but partial or absent in MHco3. MHco3 demonstrated a similar improvement with regard to CEGs that could be found as complete or partial copies in one genome, while being altogether absent in the other (Figure 1B), with 33 such cases occurring in MHco3, but only two in McMaster.

**Figure 1:**
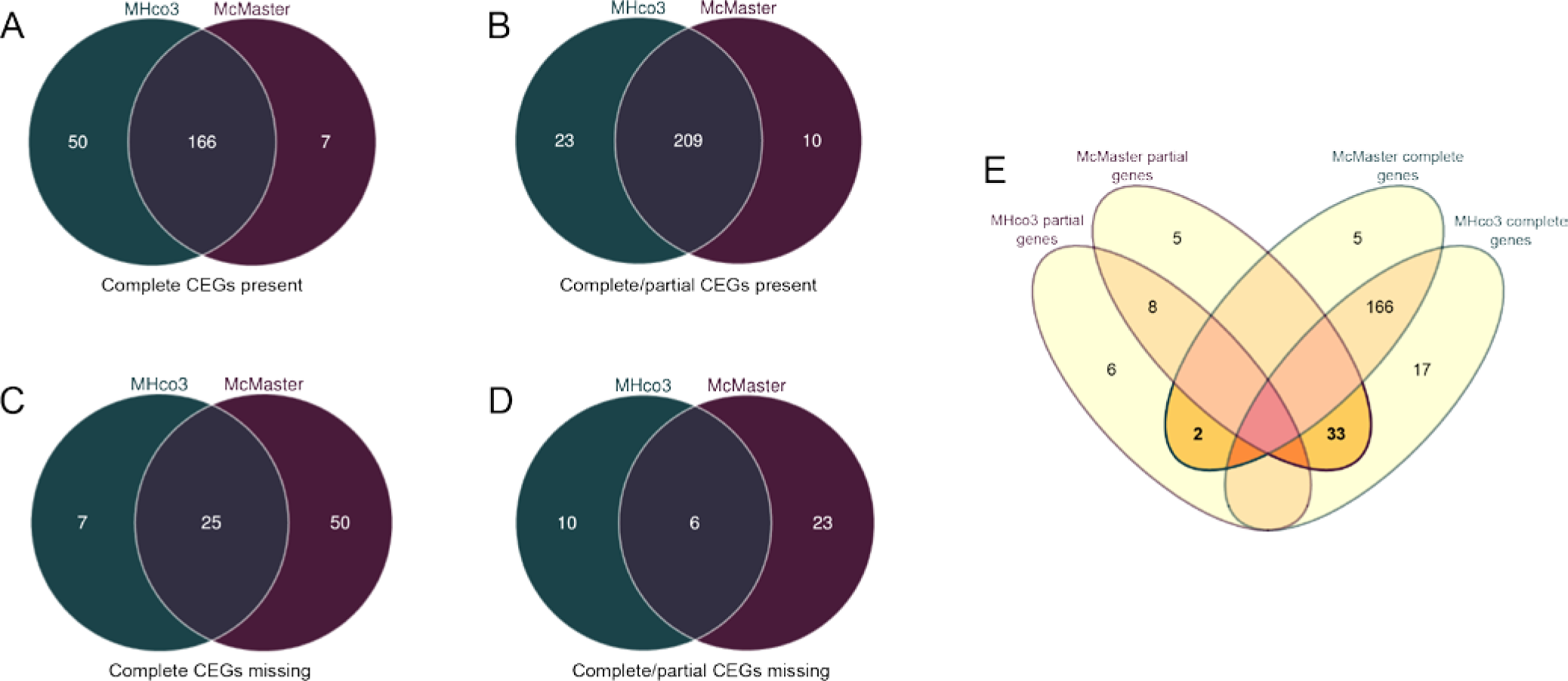
Comparison of conserved eukaryotic genes (CEGs) both present and absent across the MHco3 and McMaster genomes. These CEGs were drawn from a set of 248 genes expected to be highly conserved across eukaryotes. A complete CEG is one in which at least 70% of the corresponding sequence can be found in the genome assembly, while a partial CEG is one in which less than 70% can be found. **A.** Number of complete CEGs in each genome. Present as complete copies in both genomes are 67% (166/248) of CEGs. **B.** Number of CEGs present in either complete or partial copies in each genome. Of all CEGs, 84% (209/248) are present at least partially in both genomes. **C.** CEGs lacking complete copies in one or both genomes. These totals include CEGs that are present in partial but not complete copies. **D.** CEGs entirely missing, such that neither a complete nor partial copy can be found in its corresponding genome. **E.** CEGs complete in one genome but only partial in the other.

Turning from CEGs that were present to those that were absent, we saw a common set of 25 of 248 CEGs (10%) missing from both *H. contortus* genomes as complete copies (Figure 1C), while six (2%) could not be found even partially in either genome (Figure 1D). In considering which CEGs could not be found even partially (Figure 1D), we saw 23 missing from McMaster but not MHco3, but only 10 missing in MHco3 but not McMaster.

### Missing CEGs show clear evidence of misassembly

Of interest are CEGs present as complete copies in one genome, but only partial in the other. MHco3 again fared best (Figure 1E), with 33 complete genes in MHco3 found only as partial copies in McMaster. Conversely, only two complete genes in McMaster appeared as partial copies in MHco3.

To understand this difference, we attempted to reassemble these incompatible gene models. One example is the CEG KOG0948.4, an RNA helicase, where we saw evidence of misassembly (Figure 2). In McMaster, the first kilobase and last 600 bp of the reference gene were on one scaffold, while the middle 800 bp lay on another. Consequently, CEGMA could locate only the first portion of the gene, missing its remainder. Furthermore, portions of the reference gene were assembled out-of-order. The CEG reference fragment beginning at approximately 900 bp (Figure 2, top axis of top plot) is positioned on its McMaster scaffold before two fragments preceding it in the reference sequence.

**Figure 2:**
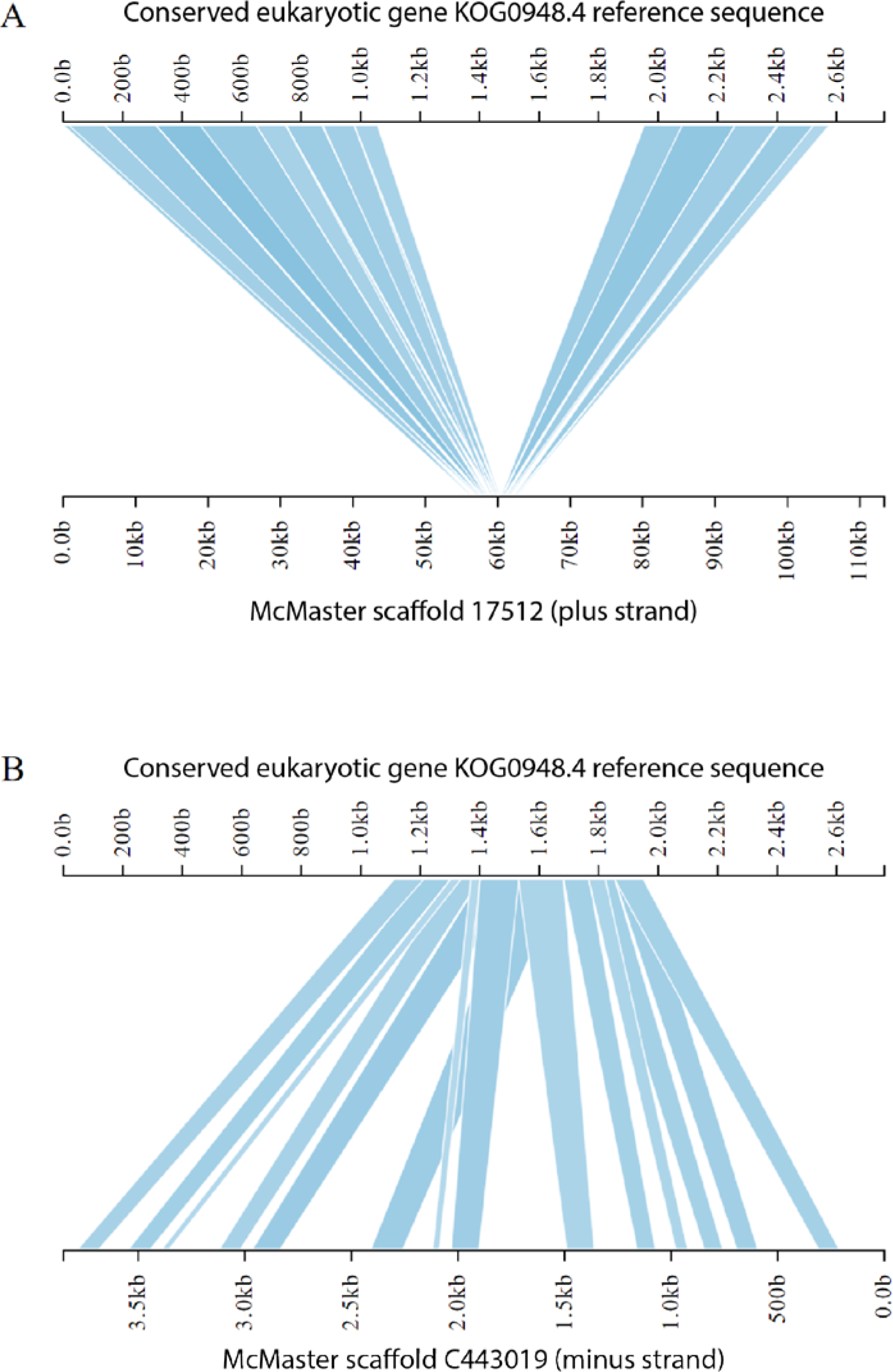
Illustration of CEG KOG0948.4 in McMaster. This CEG is present as a complete copy in MHco3 but only as a partial one in McMaster. The top axis of each plot represents the coding sequence for the CEG, while the bottom axis represents a homologous sequence found in the McMaster genome. The first 1 kb and last 600 bp of the coding sequence are found on one McMaster scaffold (top). The middle nine-hundred bases, however, are on an entirely different scaffold (bottom). Furthermore, several exons on the top plot appear on their corresponding scaffold out-of-order.

Furthermore, several such overlapping hits occur, such that a given portion of the reference sequence finds multiple hits on the corresponding McMaster scaffold.

### Both genomes poorly represent a 409 kb BAC-amplified X chromosome fragment

Seeking out a 409 kb BAC-amplified X chromosome fragment [22] in both *H. contortus* genomes demonstrated that neither is exemplary in its representation of the sequence (Figure 3). MHco3 placed the first 44 kb of the sequence on one scaffold, the region from 44 kb to 76 kb on another scaffold, and the remaining 333 kb on a third scaffold (Figure 3A). This third scaffold contained significant gaps, with as much as 23 contiguous kb missing from portions. McMaster fared worse, with the first 20 kb of the sequence placed on one scaffold, the portion from 171 kb to 246 kb on a second scaffold, and the portion from 376 to 397 kb on a third. Intervening portions of the reference fragment were scattered across myriad other scaffolds. As with MHco3, significant gaps appeared on all three scaffolds.

**Figure 3:**
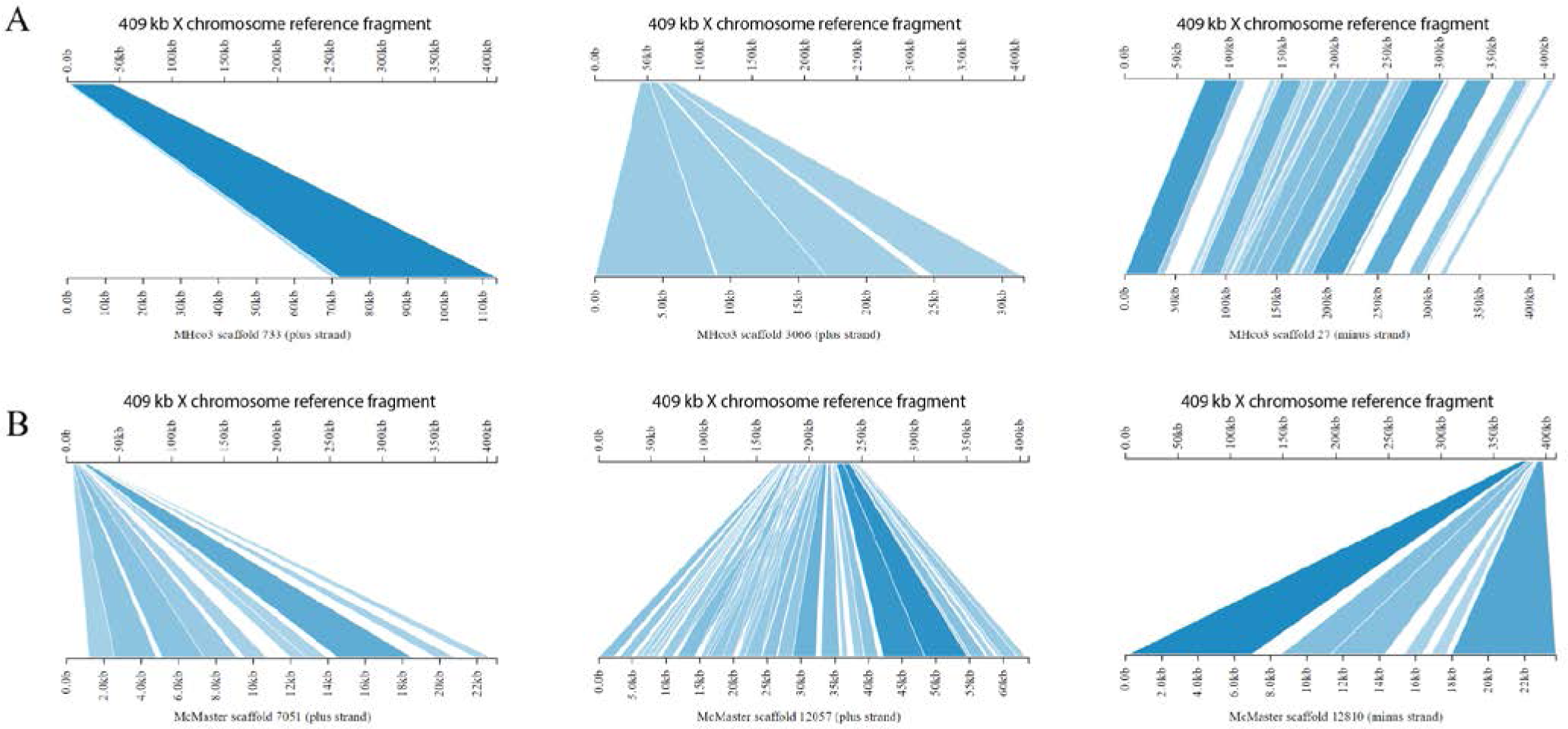
Representation of s 409 kb fragment of *H. contortus* X chromosome, sequenced from overlapping bacterial artificial chromosomes (BACs) [22]. **A.** MHco3’s representation of the 409 kb BAC. **B.** McMaster’s representation of the 409 kb BAC.

### Many gene models lack orthologues across genomes

In MHco3, only 14,539 (66%) of genes were orthologous to McMaster genes (Figure 4A). Similarly, 10,562 (45%) McMaster genes were orthologous to MHco3 genes. As both values were far lower than expected, we validated our method by computing orthology between *C. elegans* and *C. briggsae.* In this case, 74% of *C. elegans* genes found orthologues in *C. briggsae*, while 67% of *C. briggsae* genes found orthologues in *C. elegans.* These values are consistent with the respective 65% and 62% found in an earlier study [23], with our higher values likely attributable to our use of OrthoMCL rather than the earlier study’s InParanoid. We then compared each *H. contortus* genome to *C. elegans*. In MHco3, 59% of genes found orthologues in *C. elegans*, while in McMaster, 36% of genes were orthologous to ones in *C. elegans*. The reciprocal relations showed a much narrower discrepancy between *H. contortus* isolates, with 45% of *C. elegans* genes deemed orthologous to MHco3, while 42% of *C. elegans* genes were orthologous to McMaster.

**Figure 4:**
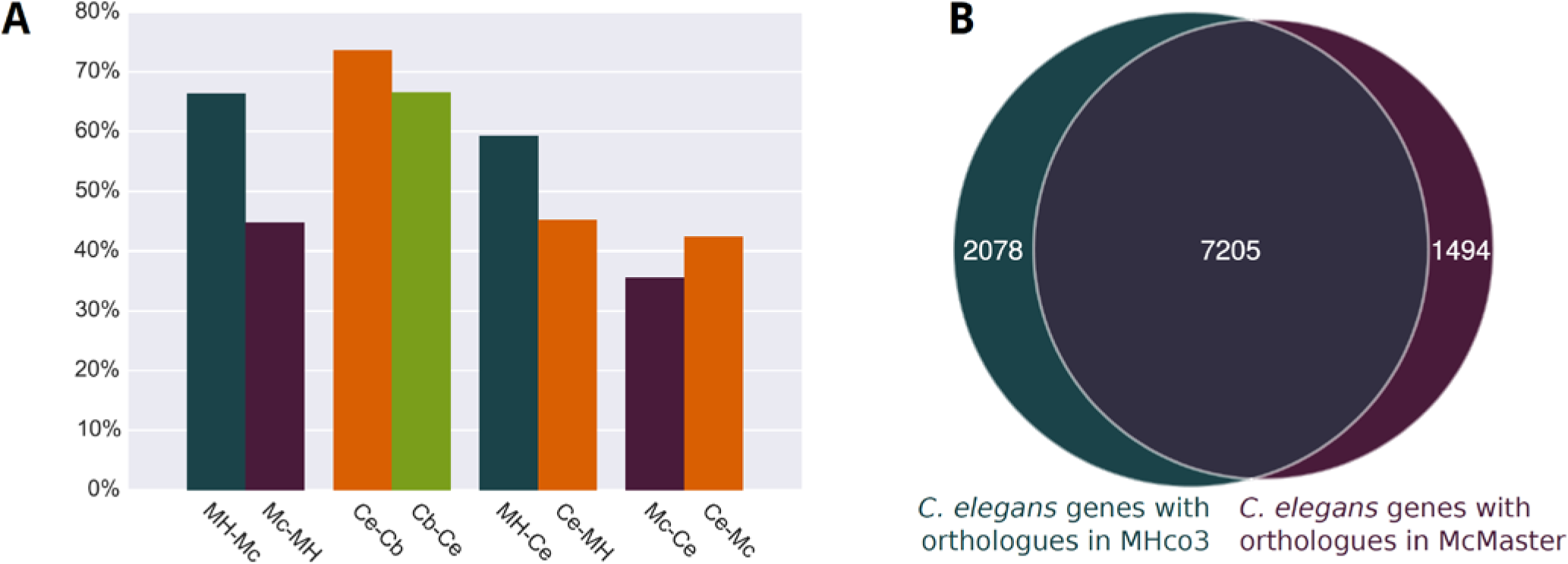
Sizes of orthologous groups of genes.. **A.** Orthologous groups between *H. contortus* MHco3 (MH), *H. contortus* McMaster (Mc), *C. elegans* (Ce), and *C. briggsae* (Cb). Note that both the forward and reciprocal relationships are depicted—e.g., MH-Mc indicates the proportion of MHco3 genes deemed orthologous to McMaster, while Mc-MH indicates the proportion of McMaster genes deemed orthologous to MH. **B.** Of *C. elegans’* 20,511 genes, 7205 (35%) were orthologous to both MHco3 and McMaster, while 2078 (10%) were orthologous to only Mhco3, and 1494 (7%) were orthologous to only McMaster.

Despite the similar magnitude of the *C. elegans* to MHco3 and *C. elegans* to McMaster orthology values, precisely which set of *C. elegans* genes were orthologous to the *H. contortus* isolates differed. Of 20,511 genes in *C. elegans,* 7205 (35%) were orthologous to both McMaster and MHco3 (Figure 4B), while 1494 (7%) were orthologous to only McMaster, and 2078 (10%) were orthologous to only MHco3.

### Missing orthologues from each genome correspond to different classifications

Though both genomes possess similar numbers of annotated proteins, each genome exhibited stark differences in the number of genes lacking orthologues in the other genome. McMaster had 13,048 genes without orthologues in MHco3 (55% of its total genes), while MHco3 had 7358 genes without orthologues in McMaster (34% of its total genes) (Figure 5A). When annotated genes without orthologues in one genome (genome A, variably referring to either MHco3 or McMaster) were aligned to annotated genes in the other (genome B), both genomes demonstrated a bimodal distribution (Figure 5B), with two-thirds of missing orthologues from each corresponding to instances in which no more than 10% of the genome A protein could be found in genome B (Figure 5C), or in which at least 90% could be located (Figure 5D).

**Figure 5:**
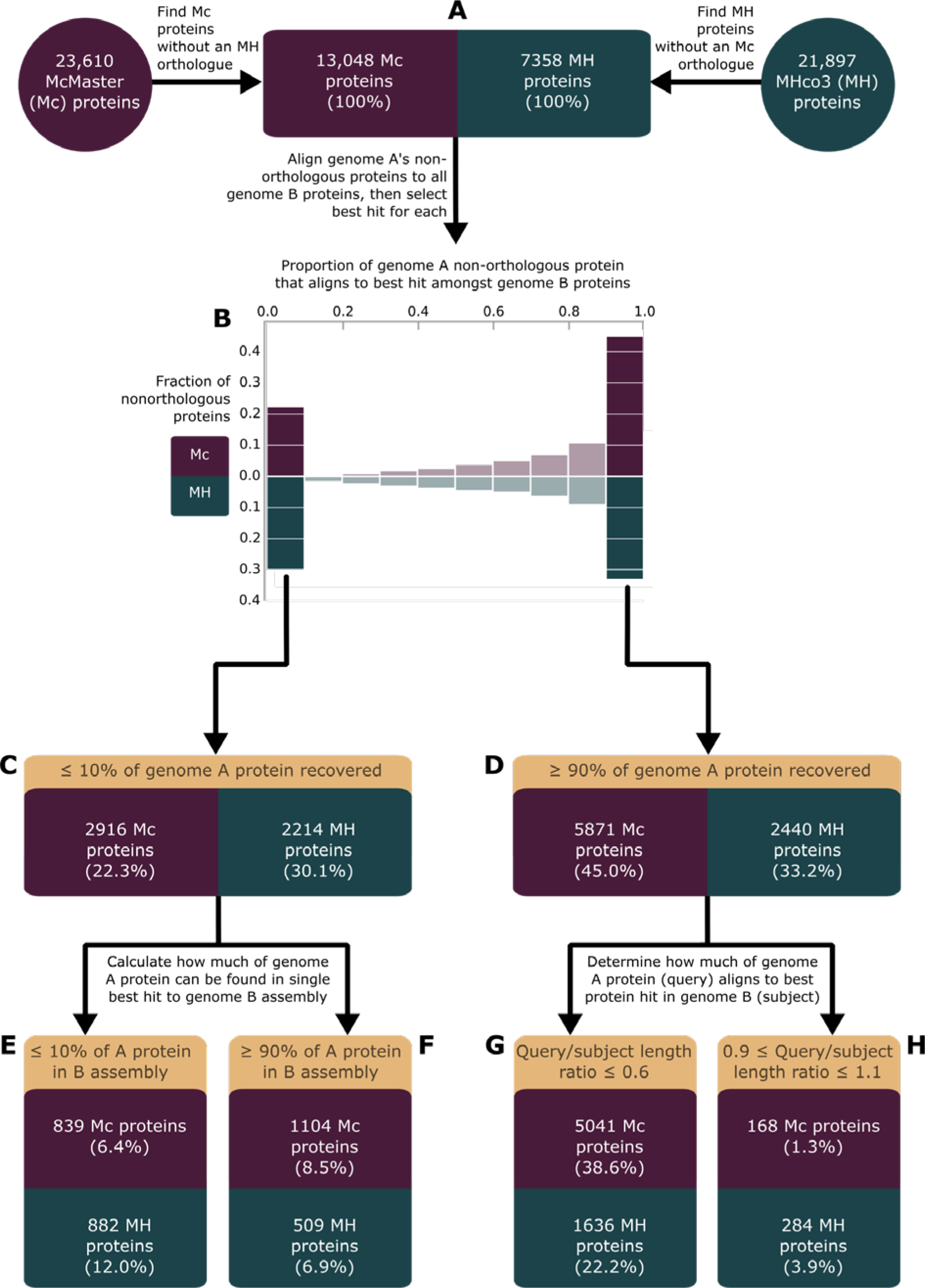
Representation of missing orthologues in each genome. Note all percentages refer to the proportion of missing orthologues in the respective genomes depicted in part A. **A.** Number of annotated proteins from each genome lacking orthologues in the other. **B.** Representation of each orthologue-less gene amongst the other genome’s gene models. **C.** Comparison of low-quality gene-model to gene-model hits between genomes. **D.** Comparison of high-quality gene-model to gene-model hits between genomes. **E.** Frequency of orthologue-less genes with low-quality hits in the other genome’s assembly. **F.** Frequency of orthologue-less genes with high-quality hits in the other genome’s assembly. **G.** Frequency of significant discrepancies between length of gene model in one genome and length of its best-hit gene model in the other genome. **H.** Frequency of similar lengths for gene model in one genome and its best-hit gene model in the other genome.

When instances in which no more than 10% of genome A’s protein could be found in an annotated gene from genome B (Figure 5C) were examined by seeking A’s protein in genome B’s assembly, a bimodal distribution against resulted, with two-thirds of these missing genes in each genome corresponding to one of two cases. In the first case, no more than 10% of the genome A protein could be found in genome B’s assembly (Figure 5E), with 6% of McMaster’s orthologue-less genes falling under this heading, and 12% of MHco3’s doing likewise. In the second case, at least 90% of the genome A protein could be found in genome B’s assembly (Figure 5. F), which occurred for 9% of McMaster’s orthologue-less genes, and 7% of MHco3’s.

The second major class of missing orthologues related to high-quality hits between gene models, in which at least 90% of the missing protein from genome A could be found in a single annotated gene in genome B (Figure 5D). Again, two cases are noteworthy. In the first case, when the length of genome A’s protein was compared to the length of genome B’s, the genome A protein was significantly shorter than the genome B one (Figure 5G)—in McMaster, 86% of such missing proteins were no more than 0.6 times the length of their best hit from MHco3, while the same was true for 67% of such proteins in MHco3. The second class of interest is that in which genome A’s orthologue-less protein was between 0.9 and 1.1 times the length of its best hit in genome B’s annotated proteins (Figure 5H). This corresponded to 1% of McMaster’s orthologue-less genes, and 4% of MHco3’s.

### MHco3 exhibits more three-way orthologues with *C. elegans*

We wished to examine synteny between the two *H. contortus* genomes, but could not identify software that adequately dealt with the large number of scaffolds in each. Consequently, we exploited the observation that while the conservation of gene order between *H. contortus* and *C. elegans* is low, genes adjacent in one species appeared together, if not consecutively, on the same chrosomome in the other species. To investigate this, we identified genes that existed as one-to-one orthologues between the two *H. contortus* genomes, as well as between each *H. contortus* genome and *C. elegans*. Such genes, thus, appeared in a 1:1:1 relationship between McMaster, MHco3, and *C. elegans,* and so are hereafter called three-way orthologues.

In total, 3037 such three-way orthologues were shared in this manner between the MHco3, McMaster, and *C. elegans* genomes. After determining the three-way-orthologue gene set, we found all scaffolds in the two *H.* contortus genomes bearing at least three three-way orthologues. Using overlapping sliding windows, we then calculated the number of *C. elegans* chromsomes onto which each contiguous triplet of three-way orthologues fell, regardless of how many non-orthologous genes intervened members of the triplet in *H. contortus*.

In McMaster, 288 overlapping windows of three contiguous three-way orthologues occurred (Figure 6A), while 1123 appeared in MHco3 (Figure 6B). Despite this four-fold difference in magnitude, the distribution of *C. elegans* chromosome counts was similar in both genomes. Each three-way orthologue group in McMaster corresponded to a mean of 1.50 *C. elegans* chromosomes (Figure 6A), while in MHco3, each three-way orthologue group corresponded to a mean of 1.48 *C. elegans* chromosomes (Figure 6B).

**Figure 6:**
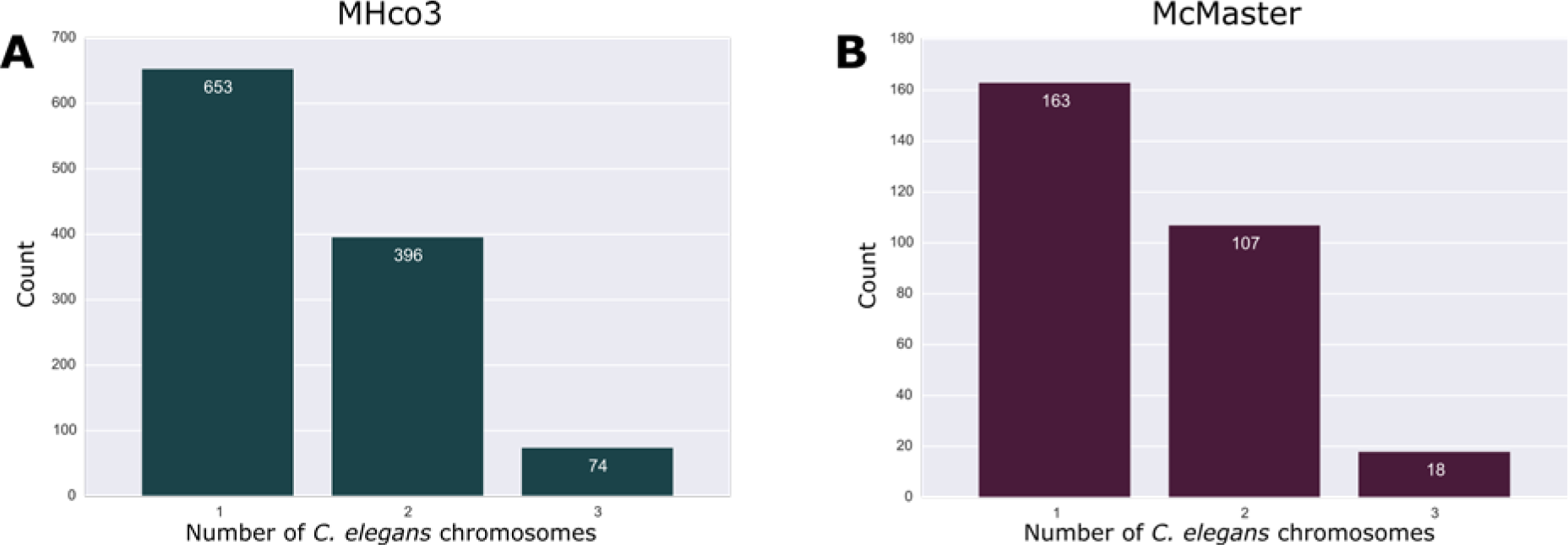
Mapping of three-way orthologue groups to *C. elegans* scaffolds. Three-way orthologues are 1:1:1 orthologues that exist in the McMaster, MHco3, and *C. elegans* genomes **A.** Each non-overlapping group of three adjacent ultraorthologues on one McMaster scaffold was classified according to whether it fell on one, two, or three *C. elegans* chromosomes. **B.** The same analysis was repeated for MHco3 ultraorthologue groups.

### Contributions to orthologous groups vary between the genomes

Of all 9159 orthologous groups established between the *H. contortus* genomes, 62.2% consisted of a single gene from each of MHco3 and McMaster (Figure 7A). Some 26.4% of the orthologous groups consisted of multiple inparalogous MHco3 genes accompanying a single McMaster gene, while only 4.9% of the groups were composed of multiple inparalogous McMaster genes alongside one MHco3 gene. Finally, 6.5% of orthologous groups included multiple genes from each genome.

**Figure 7:**
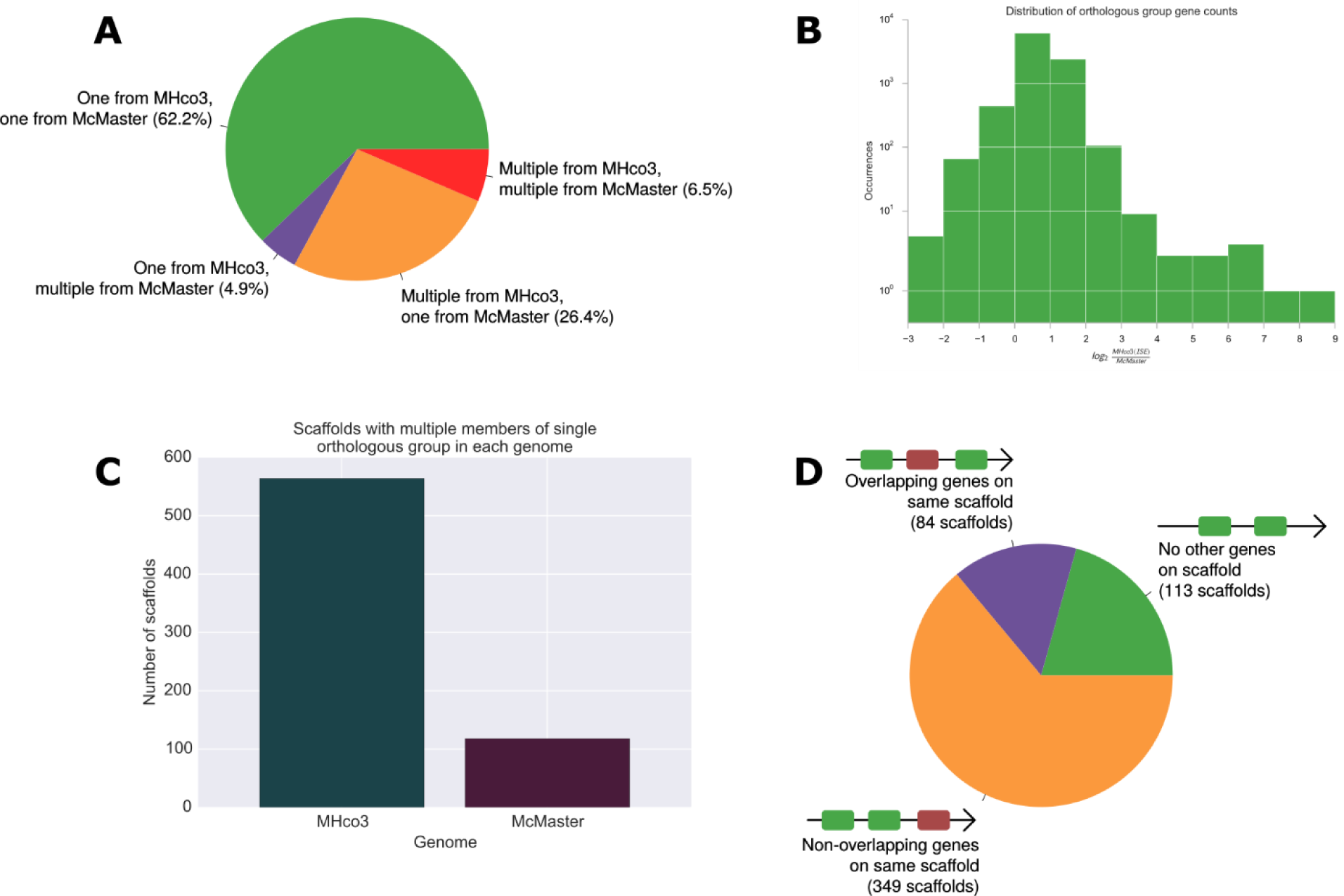
Orthologous group structure in both *H. contortus* genomes. **A.** Gene composition of 9159 orthologous groups formed between the two *H. contortus* genomes. **B.** log_2_ ratio of MHco3-to-McMaster genes in each orthologous group. The asymmetry of the x axis indicates that groups often have more MHco3 genes than McMaster genes. **C.** In MHco3, 546 scaffolds bore two or more genes from the same orthologous group, while 118 scaffolds in McMaster showed the same. **D.** Of the 546 scaffolds in Mhco3 with two or more genes from the same orthologous group, 84 have a non-orthologous gene between the orthologues, 349 have non-orthologous genes elsewhere on the scaffold but not between the orthologoues, and 113 have no non-orthologous genes on the scaffold.

Asymmetry in orthologous group structure became more apparent when we considered the precise number of genes each *H. contortus* genome contributed to the groups. Orthologous group size distribution is skewed such that groups whose MHco3 members outnumber their McMaster contingent are more common than groups exhibiting the inverse relationship (Figure 7B). Notably, there are only 463 instances in which the number of McMaster genes in an orthologous group exceeds the number of MHco3 genes by at least a factor of two, and only 17 in which there are at least four times as many McMaster genes as MHco3 ones. Conversely, there are 2517 groups in which the number of MHco3 genes surpasses the number of McMaster ones by a factor of two or more. In addition to being more numerous, such groups tend to be more dramatically skewed in favour of MHco3 genes, with 125 instances where the ratio of MHco3 to McMaster genes reaches four or more, and 18 cases where the ratio meets or exceeds eight.

### MHco3 has many more homologues placed on the same scaffolds

Consistent with the bias of multi-member orthologous groups toward MHco3 genes rather than McMaster ones, we saw 546 MHco3 scaffolds bearing multiple members of the same orthologous group, but only 118 McMaster scaffolds showing the same arrangement (Figure 7C). Of those 546 MHco3 scaffolds, 113 did not exhibit any genes outside the multi-member orthologous group (Figure 7D); 349 had at least one out-of-group gene on the same scaffold, but positioned either before or after group members such that the group remained contiguous; and 84 possessed an out-of-group gene in a position placing it between group members.

## Discussion

### Both genomes exhibit signs of misassembly

CEGMA’s set of 248 CEGs is present in most eukaryotes, spanning such diverse species as *C. elegans, Arabidopsis thaliana,* and *Homo sapiens,* and so virtually all of these CEGs will be present in each *H. contortus* isolate. CEGMA indicated, however, that MHco3 lacked 32 of 248 CEGs in complete copies, while McMaster missed 75 (Figure 1C). Given the highly conserved nature of these genes across divergent eukaryotes, this is not a biologically plausible result. As CEGMA draws solely on the genome assembly and not its associated gene models, these cases suggest misassembly. Better CEG representation in contemporary genomes is not unrealistic. For example, the carnation genome contained complete copies for 96% of CEGs [24], while the New Zealand sea urchin transcriptome represented 98% of CEGs as complete copies [25]. Both projects drew on 454 and Illumina sequencing platforms similar to those used for the two *H. contortus* genomes [7, 8].

Substantial discord appeared between the two genomes regarding which CEGs were missing. MHco3 lacked seven CEGs present completely or partially in McMaster, while McMaster missed 50 CEGs found as complete or partial copies in MHco3 (Figure 1C). Once more, this was not due to biological differences between the isolates—given the highly conserved nature of the genes, and the evolutionary proximity of the two sampled organisms, we expected almost perfect agreement between the two *Haemonchus* genomes concerning which CEGs were missing. Instances in which a CEG is present completely in one genome, but absent or only partial in the other, are likely technical errors.

Also notable is disagreement between the genomes. As CEGs are expected to appear once and only once in every eukaryote, most CEGs should not have multiple orthologues, with deviations from this guideline suggesting potential technical error. McMaster possessed a lower mean number of orthologues per CEG (Table 2), with only 1.37 orthologues found for each of its 173 complete CEGs, relative to 1.55 orthologues for each of MHco3’s 216 complete CEGs. Given that each CEG’s copy number is expected to be one, this lower value indicates that, compared to MHco3, McMaster may have better resolved CEG alleles as corresponding to the same physical locus, while MHco3 instead mistakenly assembled them as distinct genes. As MHco3 possessed a significantly larger assembly (370 Mb) than McMaster (320 Mb), with McMaster’s value falling closer to the expected true *H. contortus* genome size of 300-320 Mb (John Gilleard, personal communication), we expect that much of MHco3’s extra sequence may correspond to unresolved allelic regions.

**Table 2:** CEGMA values determined in this study. Note that these differ slightly from those published in both *H. contortus* genome papers (Table 1) despite having been calculated using the same CEGMA version.

|  | **MHco3** | **McMaster** |
| --- | --- | --- |
| CEGMA core genes (complete, of 248) | 216 (87.1%) | 173 (69.8%) |
| CEGMA core genes (total, of 248) | 232 (93.6%) | 219 (88.3%) |
| Average orthologues per CEG (complete, of 248) | 1.55 | 1.37 |
| Average orthologues per CEG (total, of 248) | 1.82 | 1.74 |

In cases where a CEG was complete in one genome but partial or absent in the other, manually seeking out portions of the CEG’s coding sequences illustrated why CEGMA failed to find the gene. Consider an example CEG complete in MHco3 but partial in McMaster (Figure 2). Several factors compromised the gene’s representation in McMaster, including the manner in which it was split across two scaffolds, the out-of-order assembly of portions of the coding sequence on one scaffold, and the seeming duplications of coding sequence such that the same fragment appeared at multiple positions on one McMaster scaffold. Furthermore, as the middle portion of the gene lay on a scaffold alone, with little accompanying sequence on the scaffold either before or after, we conclude the assembly algorithm could not properly assemble the gene as a whole, and so shunted the middle portion off to its own scaffold. Given that the coding sequence of this CEG should be extremely similar across the two *Haemonchus* isolates, these elements all suggest misassembly within McMaster.

CEG misassembly was not, of course, limited to the McMaster genome. Cases where a CEG was complete in McMaster but absent or partial in MHco3 showed the same characteristics compromising gene representation in the latter. Nevertheless, MHco3 consistently represented CEGs better than McMaster. Consider, for example, the 33 cases in which a complete MHco3 CEG was partial in McMaster (Figure 1E), compared to the mere two instances of a complete McMaster CEG being partial in MHco3. This suggests MHco3’s assembly is likely more accurate than McMaster’s—whatever factors contribute to its more comprehensive CEG representation likely benefit the assembly as a whole. It is important to note that MHco3 is an inbred lab strain, while McMaster is a field isolate [7, 8]. We would expect fewer instances of heterozygosity in the former, making it less troublesome to assemble. Furthermore, MHco3 used as its basis for sequencing a single worm, reducing instances of heterozygosity relative to McMaster, which used a sample of multiple worms. We must still, however, consider both assemblies to be flawed, for even MHco3 managed to capture only 87% of CEGs in complete copies (Figure 1A), when we expected upwards of 95% to be present *in vivo.*

Note that the CEGMA values published alongside the two *H. contortus* genomes (Table 1) differ from those determined in this study (Table 2). This was unexpected, as both publications accompanying the two genomes and our own work used the same version of CEGMA, run with its default parameters.

This discord may have resulted in part from us using more recent versions of programs on which CEGMA is dependent (such as BLAST and HMMer), or from divergences between the data set used in the genome publications and that published on WormBase. Nevertheless, these findings prompt concern about the repeatability of CEGMA analyses, given that both MHco3 and McMaster fared worse in our CEGMA runs than in their respective publications.

Apparent assembly issues extended beyond each genome’s representation of CEGs. Neither genome reproduced with complete fidelity a 409 kb BAC-amplified fragment of *H. contortus’* X chromosome [22] (Figure 3). MHco3 presented a reasonable likeness of the region, with the final 333 kb of the total 409 kb appearing on a single scaffold, albeit with significant gaps. The first 76 kb, however, were split across two other scaffolds. McMaster also faltered, with its largest single contiguous portion of the sequence corresponding to the 75 kb between 171 kb and 246 kb. The portions before and after this region were placed on multitudinous other scaffolds, again with substantial gaps. With the 409 kb BAC-amplified region serving as a high-quality representation of the *in vivo* sequence, the fragmented assembly of this region in both genomes demonstrates that assembly issues extend well beyond short regions affecting CEGs.

### Unexpectedly little orthology exists between the two genomes

We will refer to the two *H. contortus* genomes as genomes A and B to indicate that MHco3 or McMaster can stand in place of either. In genome A, almost all genes should be orthologous to one or more genes in genome B. If the two sampled isolates were identical, and the *in silico* genomes perfectly represented the *in vivo* nature of their respective isolates, we would expect a single orthologous group to be established for each gene in genome A and B, containing exactly one gene from each. Of course, given divergence of the isolates from their last common ancestor, this will not occur. Again, assuming the data sets perfectly reflect their isolates, some genes in genome A will not be placed in orthologous groups, indicating that the corresponding gene in genome B’s isolate has been deleted or diverged beyond recognition. In other cases, one gene in genome A will correspond to multiple genes in genome

B. This indicates that a duplication has occurred in genome B’s isolate, yielding multiple group members inparalogous to one another.

A comparison between two genomes for different *A. oryzae* isolates found less than 1% of genes in each were not orthologous to genes in the other [13]. Consequently, we expected in excess of 95% orthology to exist between our *Haemonchus* isolates. Even so high a value is conservative, for it permits approximately 1000 non-orthologous genes in each genome corresponding to deletion or duplication events. Whatever gap exists between the observed orthology and the expected 95% or greater value likely indicates technical error on the part of one or both genomes.

Observed orthology fell far below our expected 95% value, with only 66% of genes from MHco3 placed in orthologous groups, and even fewer—45%—from McMaster (Figure 4A). Of the 9159 orthologous groups thus formed, only 62% contained single genes from each of MHco3 and McMaster (Figure 7A), violating our expectation that the vast preponderance of groups would fall in this class.

Given these confounding results, we applied our OrthoMCL-based pipeline to *C. elegans* and *C. briggsae*, two other Clade V nematodes whose available genomic data is more refined than that published for *Haemonchus*. In comparing the two *Caenorhabditis* species to each other, we found 74% of *C. elegans* genes orthologous to *C. briggsae* (Figure 4A), and 67% of *C. briggsae* genes orthologous to *C. elegans*. These values gave us a baseline for comparison against two well-established, high-quality genome assemblies. Taken uncritically, our *H. contortus* orthology values thus suggest that two rather distant *Caenorhabditis* species that diverged 36 million years ago (MYA) [26] possess greater mutual similarity than two isolates of the same species that diverged much more recently. As this result is illogical, we question the veracity of the *H. contortus* genome data.

Next, we compared each of the two *H. contortus* genomes to *C. elegans.* In *C. elegans*, 45% of genes found orthologues in MHco3 (Figure 4A), while 42% of *C. elegans* genes found orthologues in McMaster. Of the 9283 *C. elegans* genes orthologous to MHco3, one-fifth are not orthologous to McMaster genes (Figure 4B), suggesting that though a similar number of *C. elegans* genes are orthologous to each *H. contortus* genome, there is significant discord as to precisely which *C. elegans* genes compose the orthologous sets for each.

Ignoring disagreement as to precisely which *C. elegans* genes are orthologous to *H. contortus* ones, we find that the 45% and 42% of *C. elegans* genes orthologous to MHco3 and McMaster (Figure 4A), respectively, are plausible,given the 74% orthology between the *C. elegans* and *C. briggsae* genomes. *elegans* and *C. briggsae* diverged 36 MYA [26], while *C. elegans* and *Haemonchus* last shared a common ancestor 383 MYA [27]. Of concern, however, are the reciprocal relations—59% of MHco3 genes had orthologues in *C. elegans*, while only 36% of McMaster genes found orthologues in *C. elegans*. The discrepancy between these values is problematic, for given the evolutionary proximity of the isolates, we would expect the set of genes shared by each *H. contortus* genome with *C. elegans* to be nearly identical. This divide suggests that though a similar set of genes is shared between *C. elegans* and each *H. contortus* isolate, MHco3 contains a great many more inparalogues for these *C. elegans* genes than McMaster. So large a discrepancy is biologically implausible and so suggests technical error, such that McMaster failed to assemble or annotate numerous orthologues, or MHco3 included too many. Given that MHco3 also demonstrated a greater number of average orthologues per CEG (Table 2), a class of genes whose copy number is expected to be one, and that McMaster’s smaller assembly is closer to the anticipated *in vivo* value for *H. contortus*, we conclude that MHco3’s more abundant orthologues are perhaps erroneous. In this case, MHco3 would have struggled to resolve alleles to single physical loci, instead assembling them as distinct genes appearing to be inparalogues of one another, and thus orthologues of single genes in McMaster.

### Differences in mutual orthology illustrate potential sources of technical error

To explore why genes from genome A lacked orthologues in genome B, we scored each orthologue-less genome-A gene by how much of it was present in its single best alignment to genome B’s genes. A bimodal distribution resulted (Figure 5B), with most A genes finding no more than 10% of their sequences in genome B’s annotated genes (Figure 5C), or at least 90% (Figure 5D).

For the first class, when little of genome A’s gene could be found in any genome B gene model, we sought the missing gene in genome B’s raw assembly, with two resulting cases. In the first (Figure 5E), no more than 10% of the A gene could be found in genome B’s assembly. Given the low probability that genome A both assembled and annotated the gene without any biological basis, A’s gene is likely correct, and B’s lack of it likely results from misassembly. Though this case accounts for similar absolute numbers of proteins in both genomes, when instead viewed as the proportion of orthologue-less genes in their respective genomes, it explains almost twice the share of “missing” MHco3 genes as McMaster ones. This suggests that misassmebly is more common in McMaster, supporting our CEG findings. The second case (Figure 5F), when 90% or more of the missing gene from A could be found in B’s assembly, likely stems from misannotation in one of the genomes. Either genome B failed to annotate a gene present *in vivo*, or genome A incorrectly classified as coding a region that does not serve as a gene in the organism, perhaps by mistakenly annotating a pseudogene as a gene proper.

The other major class of missing orthologues occurred when at least 90% of genome A’s orthologue-less gene could be found amongst genes annotated in genome B (Figure 5D). Most such genes from genome A had protein products much shorter than those from the corresponding annotated genes in genome B, with the A protein exhibiting a length no more than 0.6 times the length of the B protein (Figure 5G). Of all four terminal cases considered, this was by far the most common. Given a McMaster gene without an MHco3 orthologue, there was a 39% chance (corresponding to 5041 genes) that it would find a high-quailty hit in MHco3, but to a significantly longer gene; the converse relation, for MHco3 genes lacking orthologues in McMaster, explained 22% of MHco3’s orthologue-less genes, accounting for 1636 in total. This case suggests either that the genome B gene model was mistakenly extended *in silico*, perhaps by fusing two discrete genes, or that the genome A gene model was incorrectly truncated. As we expect erroneous truncations to be more common than erroneous fusions, the discrepancy between the McMaster and MHco3 values favours MHco3’s assembly and annotation.

In the final case (Figure 5H), at least 90% of genome A’s orthologue-less gene was found in a genome B gene model, and the protein-length ratios of the genome A gene and its best hit in genome B were similar, with the A gene’s length lying between 0.9 and 1.1 times that of the B gene’s. As this indicates a seemingly commensurate gene model for the genome A gene existed in genome B, these instances are the most difficult to explain. They may stem from a shortcoming in our methodolgy—as we computed the quality of each hit by determining how much of the A gene was aligned to the B gene model, without penalizing regions that overlapped or appeared out-of-order, we may have incorrectly declared a high-quality relationship between the A and B genes. *In vivo*, these cases may relate to ectopic gene conversions, in which one paralogue replaced a predecessor such that it retained high local similarity, even as regions within the gene were rearranged.

### Highly conserved Clade V nematode genes are more contiguous in MHco3 than McMaster

McMaster demonstrated 288 overlapping windows in which orthologues existing in a 1:1:1 relationship between McMaster, MHco3, and *C. elegans* appeared on the same scaffold (Figure 6A), while MHco3 exhibited 1123 (Figure 6B). This four-fold difference suggests that such three-way orthologues, consisting of highly conserved Clade V nematode genes that have undergone neither duplication nor loss, more frequently appear on scaffolds alongside other three-way orthologues in MHco3 than in McMaster.

Given the highly conserved nature of these genes, their relative chromosomal assignments are likely preserved as well. Thus, groups of three-way orthologues falling on the same scaffold in an *H. contortus* genome likely appear together on the same *C. elegans* chromosome as well. Consequently, for each triplet of three-way orthologues falling on a single scaffold in one *H. contortus* genome, we calculated whether the genes in the group fell on one, two, or three *C. elegans* chromosomes. The grouping distribution was similar for both *H. contortus* genomes (Figure 6), with McMaster’s assignments corresponding to a mean number of 1.50 *C. elegans* chromosomes per triplet, and MHco3’s groupings corresponding to a mean of 1.48 *C. elegans* chromosomes. The similarity of these distributions suggests that MHco3 did not achieve its four-fold-greater number of three-way orthologue groupings by mistakenly conjoining scaffolds. Consequently, this provides greater confidence that other indications of MHco3’s more contiguous assembly, such as its higher N50 and better X-chromosome-fragment representation, do not mislead.

### Orthologous groups contain more MHco3 genes than McMaster genes

Despite MHco3 possessing only 21,799 genes relative to McMaster’s 23,610 (Table 1), a difference exceeding 1800 genes, MHco3 contributed considerably more genes to the orthologous groups established between the two genomes—14,539 genes in MHco3 found orthologues in McMaster (Figure 4A), relative to only 10,562 in McMaster that found orthologues in MHco3. These discrepancies manifested themselves in the size of orthologous groups. Of 9159 orthologous groups, 26.4% had multiple genes from MHco3 orthologous to one from McMaster, while only 4.9% had multiple genes from McMaster orthologous to one in MHco3 (Figure 7A). A similar distinction emerged in examining the ratios of genes contributed to each orthologous group by the two *H. contortus* genomes. Of the 9159 orthologous groups, only 463 had at least twice as many McMaster genes as MHco3 ones (Figure 7B). Conversely, there were 2517 groups in which MHco3 provided at least twice as many genes as McMaster.

Taken in concert, these results illustrate that orthologous groups between the two *H. contortus* genomes consistently contained more MHco3 genes than McMaster ones, suggesting many more inparalogues exist in MHco3 than McMaster. Once more, we do not expect this divergence to be biologically valid, given the closely related nature of the sampled isolates. Consequently, we may interpret this discrepancy multiple ways, yielding different conclusions about which genome better represents inparalogous genes.

MHco3 may be the superior genome if its assembly excelled at distinguishing highly similar genes, resolving them to discrete physical loci rather than mistakenly collapsing them into single sequences. In this case, McMaster would have faltered in separating highly similar genes, a task notoriously difficult when using the short read data drawn on by both genome projects [4]. Alternatively, McMaster may be more accurate in representation of inparalogues if we conclude that MHco3 failed to resolve alleles to single genes, instead deciding that each allele corresponded to a separate locus. As previously described, McMaster’s lesser number of orthologues per CEG and closer-to-expected assembly size make this the more probable case.

### Orthologous group structure hints at biological validity

Examining the placement of orthologues provides clues as to whether apparent duplications are biologically valid. MHco3 had 546 scaffolds with at least two orthologues from the same group (Figure 7C), while McMaster had only 118. So substantial a difference favours MHco3—as the most common mechanisms of gene duplication will cause inparalogues to arise in locations physically proximal to their progenitors, the dearth of McMaster scaffolds with multiple inparalogues suggests MHco3’s assembly and/or annotation are more accurate. Though MHco3 has considerably more inparalogues than McMaster, and so will necessarily have more opportunities to place them together, the discrepancy in genes in orthologous groups (14,539 in MHco3 versus 10,562 in McMaster) is too modest to justify the observed divide.

Examining the 546 multi-orthologue scaffolds in MHco3 grants insight into their potential validity. (The proportions composing McMaster’s 118 multi-orthologue scaffolds are similar, leading to the same conclusions concerning biological validity, and so we omit their consideration.) Of the 546, 84 scaffolds had a non-orthologue positioned somewhere between genes of the orthologous group (Figure 7D). These cases are likely to represent biologically valid duplications rather than technical error—the longer and more complex an assembled region, the less likely it arose as a gross technical mistake, and so multiple genes spanning a region increases confidence in the region’s veracity. Similarly, 349 of the 546 multi-orthologue scaffolds included at least one non-orthologous gene that did not overlap the region occupied by the orthologues (Figure 7D). Again, the presence of non-orthologous genes on the scaffolds grants credence to the assembled regions’ structures. More interesting, however, are the 113 cases in which two orthologues occur on a scaffold lacking any other genes (Figure 7D), for of the three classes of multi-orthologue scaffolds, these represent the most probable instances of technical error. Tandemly arrayed repeats are difficult to assemble using short-read data [4], and so assembly algorithms may mistakenly shunt apparent repeats onto their own scaffolds when they cannot be placed in the larger genomic context. Nevertheless, we expect that tandem repeats resulting in adjacent inparalogues are the most common duplications occurring biologically, and so their existence need not be cause for concern.

### Each genome exhibits different strengths

In comparing the MHco3 and McMaster *H. contortus* genomes, several metrics indicated quality issues. These included poor representation of core eukaryotic genes and a reference X-chromosome fragment by both genomes, evidence of CEG misassembly, an unexpected lack of orthology, and discrepancies in distribution of inparalogues. MHco3 likely better represents its *H. contortus* isolate than McMaster represents its own—MHco3 demonstrated a better representation of CEGs, more instances of three-way orthologues between itself, McMaster, and *C. elegans* being assembled together, and fewer likely cases of misassembly or misannotation leading to missing orthologues relative to McMaster. Quality issues in MHco3 remain, however, as indicated by its still-wanting representation of CEGs and the X-chromosome reference sequence.

More concerning is evidence that MHco3 more frequently designated allelic regions as corresponding to distinct physical loci. Relative to McMaster, such indications include the larger average number of orthologues per CEG, an assembly size more inflated compared to the expected *in vivo H. contortus* genome, and seeming greater number of inparalogues for genes orthologous to *C. elegans*. While these issues do not compromise the aforementioned indications that MHco3 more accurately represents critical genomic regions than McMaster, the overall MHco3 assembly and annotation may also include considerably more allelic information, yielding an inaccurate illustration of *H. contortus*.

The best data set, then, depends entirely on the researcher’s aims. If the most accurate depiction of smaller-scale genomic features such as genes is required, MHco3 appears preferable; if, however, the overall representation of the organism via all assembled sequence and annotated genes is paramount, McMaster seems better. The MHco3 genome perhaps achieved its superior representation of smaller-scale genomic features because it was easier to assemble—while McMaster struggled with polymorphism in their sequenced *H. contortus* field isolate [28], MHco3 drew upon a highly inbred lab strain [7], likely reducing the number of heterozygous regions with which their assembly process had to contend. Moreover, unlike McMaster, MHco3 used single-worm sequencing data in addition to data from a multi-worm population, further reducing polymorphism. Thus, we expect MHco3’s assembly to be more accurate, which is borne out by its representation of smaller-scale genomic features. These concerns render suspect McMaster’s seemingly superior representation of global genomic structure.

McMaster may have achieved its smaller assembly by merging similar genomic regions too aggressively, under the mistaken assumption they corresponded to alleles. Such concerns are warranted given the difficulty the researchers had in assembling the McMaster genome [28], which they resolved only by discarding large volumes of sequencing data. **{Should I add a remark about how using read depth on seeming duplicated regions may help address this?}**

## Conclusions

This study presented a rare opportunity to contrast two concurrently produced genomes for the same organism. Significant discrepancies between the two genomes suggest skepticism is warranted when drawing on genomic data produced via modern sequencing platforms and assembly and annotation pipelines. While highly polished reference genomes that have undergone extensive refinement, such as those for *C. elegans* and *H. sapiens*, likely reflect well the nature of their corresponding organisms, the same may not be true for smaller projects focused on newly sequenced species such as *H. contortus*.

## Methods

All source code written for this project is published at [29]. The two *H. contortus* genomes used throughout were both drawn from WormBase WS239. MHco3 corresponds to the data set PRJEB506 [30], while McMaster corresponds to PRJNA205202 [31].

### Analyzing CEG presence

We used CEGMA 2.4.010312 [32] to determine how well the two *H. contortus* genomes represented the set of 248 highly conserved core eukaryotic genes (CEGs) included with CEGMA. In examining CEGs, “complete” CEGs are defined as those for which at least 70% of a nucleotide sequence corresponding to the CEG’s amino acid sequence could be recovered from its assembly. “Partial” CEGs are ones for which a portion of the sequence comprising less than 70% of the whole gene could be located. The CEGs used in this study were the set of 248 most-conserved CEGs selected by CEGMA’s authors.

We modified CEGMA to output the names of complete and partial CEGs, in addition to its standard summation of the number of CEGs found. For CEGs available in complete or partial copies in one genome (generically referred to as genome A) but entirely absent in the other (genome B), we extracted the coding sequence from genome A, then used BLAST 2.2.27+ [33] to search for the sequence in genome B. We repeated this BLAST procedure for genes available as complete copies in genome A but only partial copies in genome B. We then visualized BLAST results using Kablammo version dcc9833204, a web-based BLAST results viewer [34].

### Investigating representation of a 409 kb X chromosome fragment

We determined how well a 409 kb X chromosome fragment was represented by aligning it against both *H. contortus* genome assemblies using BLAST+ 2.2.28 [33]. This X chromosome fragment originated from five bacterial-artificial-chromosome (BAC) sequences bearing portions of the *H. contortus* X chromosome as an insert, sequenced and then assembled via BAC fingerprint mapping [22]. Structure of the alignments was then visualized using Kablammo version dcc9833204 [34].

### Determining orthology between genomes

To construct orthologous groups of genes between the two genomes, we used OrthoMCL 2.0.9 [35] in conjunction with BLAST+ 2.2.29 [33] and MCL 12-135 [36]. Alongside the MHco3 and McMaster *H. controtus* genomes, we used the *C. elegans* PRJNA13758 genome published in WormBase WS239 [37], and *C. briggsae* PRJNA10731 genome from WS240 [38].

Before each OrthoMCL run, we first filtered gene isoforms from the *H. contortus* data to ensure only one isoform for each gene appeared amongst the set of orthologous groups. Without this step, the size of orthologous groups would be inflated by multiple isoforms. To filter isoforms, we grouped protein sequences whose FASTA headers indicated they differed only by isoform ID, then retained the single longest isoform.

### Seeking missing orthologues

All following analyses were conducted both with genome A standing for MHco3 and B for McMaster, and also with A standing for McMaster and B for MHco3. To determine the representation of genes from each genome lacking orthologues in the other, we used BLAST+ 2.2.29’s blastp application [33] to align the annotated protein sequence of each orthologue-less gene from genome A against all annotated protein sequences in genome B. For a given gene in genome A and a corresponding hit in genome B, we computed the union of the portions of A covered by each high-scoring pair, scoring the hit-subject pairing as the ratio between the length of this union and the overall length of the A protein [39]. Using this metric, we then selected for each A gene the highest-scoring protein in genome B, defining A’s score to be the score between these two sequences. Note that the same protein in B may be assigned as the best hit for multiple proteins in A, as each comparison is conducted independently.

With each orthologue-less gene in genome A scored in this manner, we selected all A proteins with scores between 0 and 0.1, then used BLAST’s tblastn to seek a corresponding nucleotide sequence in the genome B assembly. We subsequently scored each orthologue-less A gene using the same method described above. Additionally, for all orthologue-less genes in genome A with scores ranging from 0.9 to 1.0, we computed the ratio of the length of the A protein to the length of its best-scoring hit amongst the B proteins.

## Competing interests

The authors declare that they have no competing interests.

## Authors’ contributions

JAW wrote the manuscript, which JDW continues to revise. JDW designed the project and primary analyses. JAW and GMM conducted the analyses and designed follow-up analyses.

## Acknowledgements

This work was supported by an Alberta Agriculture and Forestry grant to JDW. JAW was supported by a scholarship from Alberta Innovates.

